# Micro- and macro- heterogeneity of N-glycoproteome remodeling in endoplasmic reticulum stress

**DOI:** 10.64898/2025.12.04.692397

**Authors:** Alexander Black, Boomathi Pandi, Dominic CM Ng, Edward Lau, Maggie P. Y. Lam

## Abstract

N-glycosylation plays essential roles in the folding, trafficking, and maturation of proteins in the secretory pathways, but how individual protein and residue glycosylation rewires under endoplasmic reticulum (ER) stress is unknown. Particularly, intact glycopeptide data that retain the connectivity between glycosylation sites and the attached glycans are needed to reveal the micro- and macro- heterogeneity of N-glycosylation sites and their permutations in stressed cells. Here, we developed an optimized magnetic polyethyleneimine boronic acid-containing scaffold (mPBA) enrichment workflow to achieve sensitive and broad enrichment of intact glycoproteins for mass spectrometry analysis, quantifying 13759 unique protein-, site-, and glycoform combinations, termed glycopeptidoforms, in normal and stressed cells while requiring only 0.1 to 0.5 mg total peptide input. The data reveals a systems-level shift in the fate of hundreds of glycoproteins. N-glycosylation changes are highly dynamic, with magnitude far exceeding protein expression changes, and showing complex protein-, site-, and glycan-specific granularity. Individual glycoform reconfigurations can be observed that suggest lesions within specific steps in protein maturation and trafficking pathways. Mannose trimming is disrupted across multiple proteins and cell states, suggesting a central feature of ER stress mediated glycoproteome remodeling. Together, these results reveal molecular details into the remodeling of protein secretory pathways upon ER stress and highlight the utility of mPBA for sensitive N-glycoproteomics studies.

## Introduction

The ER is the site of biogenesis for proteins localized to the plasma membrane, cell surface, extracellular space, as well as secreted and circulating proteins. The vast majority of these proteins undergo N-glycosylation, which is essential for protein folding, stability, and trafficking along the secretory pathway (1, 2). In mammals, the attachment of mature N-linked glycosylation onto proteins is an orchestrated, multi-step process involving hundreds of enzymes (3, 4). The protein portion of the process begins with the transfer of GlcNAc_2_Man_9_Glc_3_ to the asparagine of the N-X-S/T sequon in proteins, either co-translationally to the nascent polypeptide chain by OST-A, or post-translationally via OST-B in the ER lumen (5). This is rapidly followed by the removal of 2 glucoses (Glc) by glucosidases I and II; the resulting monoglucosylated attached protein is recognized by the two lectin chaperones calnexin (CNX) and calreticulin (CRT) to assist in protein folding and further deglucosylated to GlcNA_c_2Man_9._ Successfully folded glycoproteins are anterograde trafficked to the Golgi and undergo trimming of mannose residues from GlcNAc_2_Man_9_ to GlcNAc_2_Man_5_, catalyzed by the Golgi ɑ-mannosidases MAN1A1, MAN1A2, and MAN1C1. Glycoproteins that fail to fold properly in the ER may be re-glucosylated by UDP-glucose glycoprotein glucosyltransferase 1 (UGGT1) to undergo additional folding cycles, or subjected to ER mannose trimming by ER degradation-enhancing alpha-mannosidase-like 1 and 3 (EDEM1/3) to be marked for proteolysis by ER-assisted degradation (ERAD). Thus, protein N-glycosylation not only allows proteins access to a highly regulated folding and proteostasis pathway, but also serves as a molecular tag that reflects the maturation and trafficking status of the modified protein. Consistent with its key role in producing and maintaining the cell surface proteome, the disruption of the N-glycosylation pattern at specific proteins is sufficient to drive large shifts cellular phenotypes ranging from membrane ion conductance to drug sensitivity and cancer immune evasion (6–8).

ER stress is a central feature in multiple diseases including cancer, neurodegeneration, and cardiac hypertrophy. Under stress conditions including oxidative stress and gene mutations, the capacity of protein folding in the ER becomes overwhelmed, leading to a backlog of protein folding, glycosylation defects, and accumulation of misfolded proteins (9, 10). ER stress conditions trigger the unfolded protein response (UPR) pathway, consisting of three main branches (PERK, IRE1, ATF6). UPR restores proteostasis by suppressing global protein synthesis while inducing stress response genes that function in part by modulating glycosylation, including IRE1/XBP1-mediateed stimulation of hexosamine biosynthesis (11) and differential expression of glycoenzymes (12). Recent work has begun to reveal the systematic effects of ER stress and UPR on protein N-glycosylation. In glycomics experiments, Wong et al. showed that under thapsigargin-induced UPR, cell membrane protein attached glycans show an increase in high-mannose classes and reduced sialylation in a cell-type-dependent manner (12). In parallel, Xu et al. (13) used bio-orthogonal N-azidocetylgalactosamine (GalNAz) labeling, click chemistry enrichment, and Peptide:N-glycosidase F (PNGase F) deglycosylation to identify 981 deglycosylated NXS/T sites after chemical ER stress induction, revealing a global alteration of glycosylation site occupancy. Taken together, these prior studies provide evidence that the glycoproteome undergoes substantial remodeling under ER stress conditions. However, the connectivity of which individual glycan motifs are attached to each protein and each N sequon, and how these configurations permute under stress, remain unknown.

Substantial variability exists in the configuration of N-glycoproteins, where changes in N-glycosylation profiles may involve: (1) the overall glycoprotein abundance; (2) each NXS/T glycosylation site on the glycoproteins being differentially glycosylated (macro-heterogeneity); and (3) different N-glycan structures at one site (micro-heterogeneity). Taken together, each pool of the same N-glycoprotein primary sequence can have distinct combinations of glycans on each sequon, leading to substantial glycoproteoform complexity (“meta-heterogeneity”) (14). Although glycomics and “deglycoproteomics” experiments where proteins are artificially deglycosylated can both offer partial information on glycosylation status, revealing the full gamut of macro- and micro- heterogeneity requires glycoproteomics approaches where intact glycopeptides are preserved to allow for site-specific as well as site- and glycoform-specific analysis (15–17). Advances in mass spectrometry data acquisition and software automation now enable the simultaneous fragmentation and assignment of peptide and glycan ions in intact glycopeptide analysis using accessible high-energy collision induced dissociation (HCD) fragmentation mass spectrometry (18–21). However, intact glycopeptides analysis remains challenging due to their poor ionization, complex fragmentation patterns, and low abundance (17, 22). These challenges have hindered intact glycopeptide comparisons across cell states and necessitate the continual refinement of glycopeptide enrichment strategies (22, 23).

In this study, we systematically investigated how ER stress influences N-glycosylation patterns in stressed human cells. To enrich glycopeptides for mass spectrometry analysis, we implemented and optimized a magnetic polyethyleneimine boronic acid-containing scaffold (mPBA) method. The combination of polyethylenimine dendrimer mesh with boronic acid chemistry allows improved glycocapture efficiency, whereas conjugation to magnetic beads streamlines processing and facilitates automatability potential such as using robotic-assisted protocols (24, 25). Comparisons of incubation pH, organic, and ionic strengths revealed conditions conducive to capturing multiple classes of N-glycans on glycopeptides. Coupled to HCD fragmentation mass spectrometry, this method allowed the quantification thousands of endogenous glycopeptidoforms across from low sample input, including 10189 from 0.1 – 0.5 mg HeLa cells; 4129 in AC16 cells at 0.5 mg input per sample; and 4777 from 0.2 mg human induced pluripotent stem cell-derived caridomyocytes (hiPSC-CM). Applying this workflow to chemical ER stress induction via thapsigargin and tunicamycin, we find that ER stress provokes a pervasive remodeling of N-glycosylation in a protein-, site-, and glycan-specific manner, broadly influencing ER resident, cell surface, secreted, and UPR-induced proteins with implications on protein trafficking and homeostasis. The analysis further reveals a proteome-wide increase in abundance of high-mannose glycopeptides and an accompanying decrease in trimmed mannose species, indicating defective mannose trimming is a common mechanism in ER stress-mediated glycoproteome remodeling.

## Methods

### Derivatization of magnetic polyethyleneimine boronic acid-containing scaffolds

To generate the magnetic polyethyleneimine boronic acid-containing scaffold (mPBA), we aliquoted 1 mL of MagnaBind Carboxyl derivatized beads which corresponds to 20 mg of bead material, or 4.8 μmol of carboxylic acid. The bead solution was then removed using a magnetic rack. The beads were then washed three times with 1 mL of PBS before resuspension in a 1.5 mL solution of 0.1 M MES buffer pH 6.0 containing 13 mg of DMTMM. This solution was mixed for 5 minutes with end-over-end mixing. 300 mg of branched PEI MW:25000 was added to the solution and mixed for 24 hours of end-over-end mixing. After PEI addition, the supernatant was removed, and the beads were washed with 1 mL of methanol 3 times. Finally, the beads were resuspended in 1.86 mmol of 5-carboxybenzoboroxole and 2.88 mmol of DMTMM dissolved in 20 mL methanol. The reaction was allowed to continue for 24 hours with end-over-end mixing. Beads were then diluted 2× with methanol and stored at 4°C.

### Human cell culture and hiPSC-cardiomyocyte differentiation

Three human cell lines were used in this study including HeLa cells, AC16 (cardiomyocyte-fibroblast hybrid), and hiPSC-derived cardiomyocytes (AICS-52). HeLa (passage 6) and AC16 cells (passage 7) were cultured in DMEM/F-12 media supplemented with 10% and 1% FBS (v/v) respectively. For the ER stress samples, passage 7 AC16 cells were grown to 70% confluency then treated with either 1 μM of thapsigargin or 1 μg/mL of tunicamycin or vehicle (DMSO) for 24 hours.

hiPSC (AICS-52 from WTC-11 background) was acquired from the Allen Cell Collection via Corielle. Cells were maintained on GelTrex-coated plates with StemFlex media supplemented with mTeSr Plus (STEMCELL). 80% Confluent hiPSC were differentiated into cardiomyocytes by culturing in RPMI 1640 + B27-insulin + 6 μM CHIR for 2 days. Afterwards CHIR was removed for 1 day. Then, RPMI 1640 + B27 -insulin and 5 μM of IWR was added for 2 days before IWR removal for another two days. Finally, RPMI 1640 + B27 + insulin was added until spontaneous hiPSC-CM contraction was observed. The hiPSC-CM were cultured in RPMI 1640 + B27 + insulin + 2 μM CHIR until 60% confluent. CHIR was removed for 13 days prior to treatment with either 1 μM of thapsigargin or 1 μg/mL of tunicamycin or vehicle (DMSO) for 24 to 72 hours. All cells were stored in a humidified atmosphere containing 5% CO_2_ at 37 °C in an incubator.

### Cell lysis, protein extraction, and protein digestion

AC16 or HeLa cells were harvested by first removing media and then washing with 10 mL of sterile PBS without Ca^2+^ or Mg^2+^. Next cells were dissociated using 2mL of 0.25% trypsin-EDTA was added to each T-75 flask and incubated at 37 °C for 2-3 minutes. Trypsin was then neutralized with 8 mL of DMEM/F-12 media supplemented with 10% FBS. Afterwards, the cells were centrifuged and washed with PBS, then snap-frozen in liquid nitrogen and stored at -80 °C until ready for further processing. Once ready, frozen cell pellets were dissolved in RIPA buffer + HALT protease and phosphatase inhibitor. The lysate was sonicated with the Biorupter Pico system for 10 cycles of 30 seconds on, 30 seconds off. After sonication, the lysate was centrifuged at 14,000 x G for 5 minutes at 4 °C to remove any cell debris. Protein concentration was then determined using the Qubit protein broad range kit. Cells in the hiPSC-CM experiments were harvested by first removing media and then washing with 10 mL of sterile PBS without Ca^2+^ or Mg^2+^. 3 mL of Triple select 10× was added to each T-75 flask and incubated at 37 °C for 15 minutes. Cells were diluted in RPMI 1640 before centrifugation and then washed with PBS, then snap-frozen in liquid nitrogen and stored at -80 °C until ready for further processing. Protein extraction followed the same format as HeLa and AC16 cells.

All samples were subjected to SP4 bead-free cleanup prior to trypsin digestion (26). A total of 0.1 – 1 mg of HeLa protein was aliquoted into new 2 mL Eppendorf protein lo-bind tubes. A total of 0.5 mg protein was used for AC16 samples, and 0.2 mg for hiPSC-CM samples. Samples were then reduced with 10 mM DTT for 30 minutes at 56 °C and alkylated with 15 mM iodoacetamide for 30 minutes in the dark. Next, acetonitrile was added to 80% v/v and the samples were briefly vortexed and centrifuged at 15,000 ×g for 5 minutes. Protein pellets were washed three times with 80% ethanol and centrifuged at 15,000 ×g for 2 minutes. Protein pellets were then resuspended in 0.3 mL of 50 mM ammonium bicarbonate and sonicated using a sonicator probe (20% amplitude 20 seconds). Mass spectrometry grade trypsin was added at a 1:100 enzyme:protein ratio overnight at 37 °C.

### Selective glycopeptide capture and elution using mPBA scaffolds

For each sample, an aliquot of mPBA beads equivalent to 25% of total peptide mass was used. The PBA bead storage buffer (methanol) was removed with a magnetic rack prior to glycopeptide enrichment. The beads were rinsed with 200 µL of MeOH, followed by 170 µL of 1% TFA in 50% ACN after removal of the MeOH. To equilibrate the beads, 200 µL of loading buffer (50 mM sodium carbonate in 60% ACN, pH 10.5) were added to each sample tube. This step was repeated once after the previous loading buffer was discarded. Peptides in loading buffer (1:1.5 peptide to buffer ratio, w/v) were then added onto the beads and allowed to incubate at room temperature for 15 minutes in a ThermoMixer at 1,000 rpm. Following incubation, the supernatant containing the non-glycosylated peptides was transferred to new eppendorf tubes. Beads were rinsed 3 times using the loading buffer (150 µL of buffer per 50 µg of beads), and glycopeptides were then eluted from the beads after incubation in elution buffer (100 mM Fructose in 1% TFA, 50% ACN, at 75 µL per 100 µg peptides) in a ThermoMixer at room temperature for 30 minutes at 1,000 rpm. The eluted glycopeptides (sample-G) and the non-gycosylated peptides (sample-N) were dried with a speed-vac and cleaned up by C18 spin columns (Thermo #89873).

### Intact glycopeptide analysis by mass spectrometry

Extracted glycopeptide samples were resuspended in 0.1% formic acid in 3% MS-grade acetonitrile and transferred into LC-MS autosampler vials. For glycopeptide samples, the entire sample volume was injected via a Vanquish Neo UHPLC system (Thermo Fisher Scientific) to an Orbitrap Exploris 480 (Thermo Fisher Scientific) mass spectrometer. Glycopeptides were separated on an EasySpray^TM^ C18 analytical column (PepMap C18, 3 μm, 100 Å, 75 μm x 15 cm) heated to 50 °C, over a 120-minute (hiPSC-CM ER stress and HeLa curve) or 90-minute (Optimization experiments) gradient with a flow rate of 200 nL/min. The mobile phase consisted of solvent A (0.1% FA in 100% LCMS-grade water) and solvent B (0.1% FA in 80% LC-MS-grade ACN) with a 90 minute-gradient of solvent B from 1–23% for 78.2 minutes, 23–50% for 5 minutes, 50–99% for 1.5 minutes, and a hold at 99% for 4.6 minutes or a 120-minute gradient of solvent B from 1–23% for 104.3 minutes, 23–50% for 10.2 minutes, 50–99% for 1.3 minutes, and a hold at 99% for 4 minutes. The mass spectrometer operated in positive mode with MS1 scan acquired from 505-2000 m/z at 60,000 resolution in profile mode using a maximum injection time of 60 seconds with a normalized AGC target of 300% and RF lens of 50%.

For the mPBA condition optimization experiments, glycopeptide precursor ions were selected following a data-dependent acquisition with a TopN of 11. Ions with charge states 2-7 and intensity greater than 3.0E4 were selected for fragmentation using a 2 m/z isolation width and accumulated for up to 240 ms with the normalized AGC target set to 500% and fragmented using stepped-HCD at 22-32-42 NCE. MS2 scans were acquired with a fixed first mass set to 100 m/z at 90,000 resolutions in profile mode. Selected ions were then placed on a dynamic exclusion list for 45 seconds. For HeLa concentration curve and hiPSC-CM ER stress experiments, glycopeptide precursor ions were selected following a data-dependent acquisition with a TopN of25. Ions with charge states 2–7 and intensity greater than 3.0E4 were selected for fragmentation using a 2 m/z isolation width and accumulated for up to 90 ms with the normalized AGC target set to 400% and fragmented using stepped-HCD at 22/32/42 NCE. MS2 scans were acquired with a fixed first mass set to 100 m/z at 45,000 resolution in profile mode. Selected ions were then placed on a dynamic exclusion list for 100 seconds.

For the AC16 ER stress experiments, all the resuspended glycopeptide samples were injected into an EasynLC-1200 (Thermo Fisher Scientific) coupled to a Q-Exactive HF (Thermo Fisher Scientific) mass spectrometer. Glycopeptides were first trapped on a C18 trapping cartridge (Acclaim PepMap^TM^ 75 μm × 2 cm) then separated on an EasySpray^TM^ C18 analytical column (PepMap C18, 3 μm, 100 Å, 75 μm × 15 cm) heated to 50 °C, over a 120-minute gradient with a flow rate of 300 nL/min. The mobile phase consisted of solvent A (0.1% FA in 100% LCMS-grade water) and solvent B (0.1% FA in 80% LCMS-grade ACN) with a 90 minute-gradient of solvent B from 2–26% for 100 minutes, 26-99% for 13 minutes, and a hold at 99% for 7 minutes. The mass spectrometer operated in positive mode with MS1 scan acquired from 500-2000 m/z at 60,000 resolution in profile mode using a maximum injection time of 70 seconds with an AGC target of 3e6. Glycopeptide precursor ions were selected following a data-dependent acquisition with a TopN of 8. Ions with charge states 2–7 and intensity greater than 1.0E4 were selected for fragmentation using a 2 m/z isolation width and accumulated for up to 240 ms with the AGC target set 5e5 and fragmented using stepped-HCD at 20-30-40 NCE. MS2 scans were acquired with a fixed first mass set to 100 m/z at 60,000 resolution in profile mode. Selected ions were then placed on a dynamic exclusion list for 60 seconds.

### Mass spectrometry for total proteome analysis

For AC16 ER stress total proteome analysis experiments, 200 ng of peptide digest was injected via a Vanquish Neo UHPLC system (Thermo Fisher Scientific) to an Orbitrap Exploris 480 (Thermo Fisher Scientific) mass spectrometer. Peptides were separated on an EasySpray^TM^ C18 analytical column (PepMap C18, 3 μm, 100Å, 75 μm x 15 cm) heated to 50 °C, over a 180-minute gradient with a flow rate of 200 nL/min. The mobile phase consisted of solvent A (0.1% FA in 100% LCMS-grade water) and solvent B (0.1% formic acid in 80% LC-MS-grade acetonitrile) gradient of solvent B from 1-20% for 114 minutes, 20-30% for 41 minutes, 30-67% for 18 minutes, 67-99% for 0.5 minutes, and a hold at 99% for 5 minutes. The mass spectrometer operated in positive mode with MS1 scan acquired from 400–1000 m/z at 60,000 resolution in profile mode using a maximum injection time set to auto with a normalized AGC target of 300% and RF lens of 50%. Peptide precursor ions were selected following data-independent acquisition with 6 m/z isolation windows from 400-1000 m/z with optimized placement on. The maximum ion accumulation time was set to auto and the normalized AGC target set 800% and fragmented by HCD at 30 NCE. MS2 scans were acquired with a fixed first mass set to 100 m/z at 22,500 resolution in profile mode with loop control set to all.

### Database search, data analysis, and statistics

All intact-glycopeptide data were searched using FragPipe v.23.0 (27) with MSFragger v.4.2 and IonQuant v.1.11.9 (28). A modified version of the glyco-N-LFQ workflow was used and the MS spectra was searched against UniProt Swiss-Prot (29) canonical human protein database (downloaded December 2023), with bovine serum albumin and ovalbumin appended. The search parameters were as follows: Peptide length between 7–50 residues, up to 2 missed cleavages allowed, precursor mass tolerance 20 ppm, fragment mass tolerance at 20 ppm, maximum three variable mods (including oxidation of methionine, protein N-term acetylation). Carbamidomethylation of cysteine was assigned as a fixed modification. The N-glycans were assigned with the Human_N-glycans_large database with all HexNAc(0) entries removed, using PTM-Shepherd in FragPipe, requiring 1% peptide decoy FDR for confident identification (30). Phosphorylation of one hexose was enabled as a variable glycan modification. MS1 label-free quantification was performed using IonQuant v1.11.9 (28) with match-between-runs enabled.

For total protein abundance analysis experiments, AC16 data acquired with DIA were searched using DIA-NN v2.0 (31). The MS spectra was searched against UniProt Swiss-Prot canonical human protein database with bovine serum albumin and ovalbumin appended (downloaded Decemeber 2023). An in-silico spectral library was generated from this fasta file for the search. The search parameters were as follows for AC16 cells: peptide length between 6 and 48 residues, precursor charge between +1 and +5, up to 2 missed cleavage allowed. Up to 2 variable modifications including oxidation of methionine and protein N-terminal acetylation were allowed and carbamidomethylation of cysteine was assigned as a fixed mod. Match-between-runs was enabled and proteotypicity was based on protein names of the fasta file. Protein total abundance data from hiPSC-CM samples were searched with identical parameters with the following modifications: peptide length between 6 and 45 residues; precursor charge between +2 and +4, up to 1 missed cleavage allowed; variable methionine oxidation.

Protein and glycopeptidoform comparisons were performed using the DEP2 package (32) in R. Unless otherwise stated, an FDR adjusted P value of 0.05 or below is considered significant.

## Results

### mPBA enables efficient capture and release of glycopeptide classes from low input

To compare the global intact glycopeptide profile of different cellular conditions, we first sought to implement a sensitive glycopeptide enrichment protocol based on boronic acid chemistry. Briefly, boronic acids reversibly forms cyclic esters with cis-diols such as found in multiple types of sugars (mannose and sialic acids) in a pH-dependent manner (**Figure 1A**). Diol capture occurs when pH is above the pKa of conjugated boronic acids, whereas release occurs when pH is below pKa. This reversibility allows reactions to flexibly capture and wash off glycans under relatively mild conditions. A hindrance to the use of boronic acid has been its relatively weak glycan binding affinity, but this may be overcome by conjugating multiple boronic acid derivatives to dendrimers to enhance overall enrichment avidity (33–35). To implement a multivalent boronic acid-based workflow, we therefore conjugated magnetic beads to branched polyethylenimine (PEI) overnight, followed by derivatization of benzoboroxole. This schema allows ease of processing for comparing binding conditions and future automatability through the use of magnetic beads, whereas the multiple valencies of glycan capture on dendrimerized boronic acids increases capture efficiency (**Figure 1B**). The resulting mPBA scaffold is used to bind to glycans from a complex peptide mixture (**Figure 1C; STEP 1**). Upon removal of unbound peptides in the wash step (**Figure 1C, STEP 2**), glycan binding is reversed to elute glycopeptides for mass spectrometry analysis (**Figure 1C; STEP 3**).

**Figure 1:**
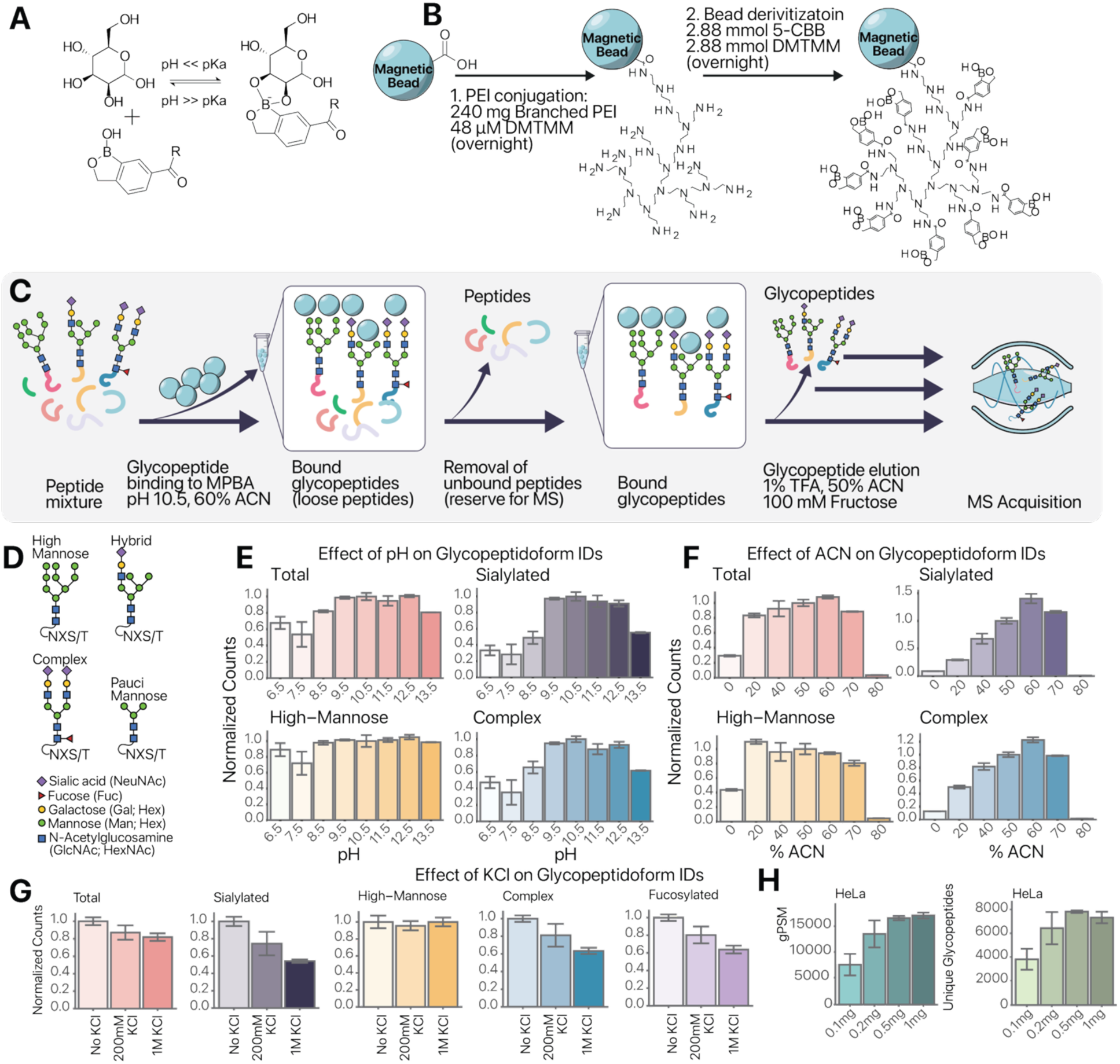
mPBA enables efficient N-glycopeptide enrichment with different glycan types. **A:** Schematic of the reversible reaction of benzoboroxole with a cis-diol containing sugar (D-mannose). Covalent bond formation is favored at high but not low pH. **B:** mPBA derivatization procedure. Step 1: branched polyethyleneimine (PEI) dendrimer is crosslinked to the carboxylic acid groups on magnetic beads. Step 2: benzoboroxole is crosslinked to primary amines on the PEI scaffold **C:** Intact glycopeptide enrichment workflow. Step 1: Incubation of protein digest containing glycosylated and non-glycosylated peptides with mPBA scaffold. Step 2: Unbound peptides are removed at the wash step. Step 3: Bound glycopeptides are eluted for mass spectrometry analysis. **D:** Common classes of N-glycans in mammalian cells including high-mannose, hybrid, and complex types. **E:** Effect of binding buffer pH has on the identification rate for total glycopeptides, high mannose containing glycopeptides, complex glycopeptide and sialylated glycopeptides (n=2 replicates × 8 conditions). **F:** Effect of binding buffer acetonitrile (ACN) concentration has on the identification rate for total glycopeptides, high mannose containing glycopeptides, complex glycopeptide and sialylated glycopeptides (n=2 replicates × 6 conditions). **G:** Effect of binding buffer KCl concentration has on the identification rate for total glycopeptides, high mannose containing glycopeptides, complex glycopeptide and sialylated glycopeptides (n=2 replicated × 3 conditions). **H:** Number of total glycopeptide-spectrum matches (gPSM) and distinct glycopeptidoforms identified in HeLa cells using the optimized mPBA protocol, at different masses of starting protein material (n=2 replicates × 4 condition).

Most human N-glycans possess the same penta-saccharide Man_3_GlcNAc_2_ core, from which extensions distinguish the high-mannose N-glycans from hybrid glycans and complex type glycans that contain fucosylated and sialylated sugars. In rare cases, paucimannose glycans with further trimmed scaffold may also occur especially in the lysosome and in tumors (**Figure 1D**). We therefore evaluated the efficacy of the mPBA scaffold towards different types of common human glycosylation structures. To optimize the efficacy for total glycopeptide enrichment as well as the relative enrichment of different glycopeptide classes, we compared the effects of pH, organic strength and salt concentration during the binding and elution steps, as these parameters have been cited to influence boronic acid chemistry for total glycopeptide identification (35, 36). We found that mPBA is compatible with a relatively wide range of basic pH from 9.5 to 12.5 for total glycopeptide and high-mannose glycopeptide enrichment, but the performance on complex and sialylated glycopeptides is highest at pH 10.5, though the effect is relatively minor, suggesting multiple pH conditions may be suitable (**Figure 1E**). On the other hand, acetonitrile (ACN) percentage has contrasting effects on the different classes of glycopeptides. Whereas the enrichment for total glycopeptides is broadly effective from 20% to 70% acetonitrile, peaking at 60%, low-acetonitrile conditions (e.g., 20%) show a relative preference for the capture of high-mannose glycopeptides. There is a substantial (>20%) gain in the efficacy of capturing complex and sialylated glycopeptides at 60% acetonitrile compared to 50% (**Figure 2E**). Thirdly, we tested the effect of salt addition during the binding phase and found that no amount of KCl led to an improvement in enrichment but rather had severe detrimental effects on the enrichment of sialylated and complex glycans (**Figure 2F**). Taken together, we conclude that a moderate organic strength, of 50% or 60% ACN, is generally suitable for glycopeptide enrichment, but may bias toward different glycan classes in different experiments, whereas contrary to prior observations (35), the addition of KCl has no benefits in our setup.

**Figure 2:**
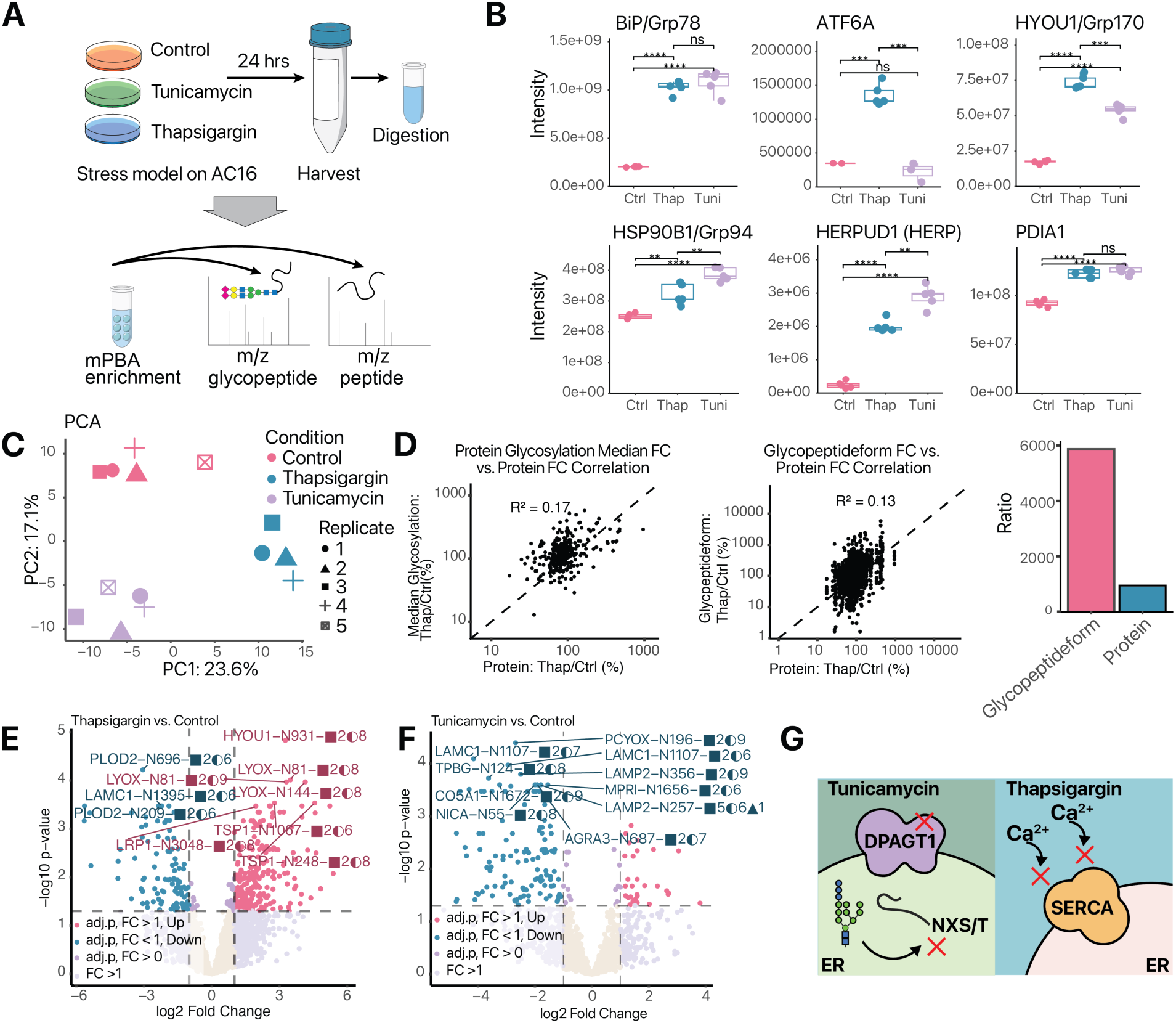
ER stress induces broad glycoproteome remodeling beyond proteome changes. **A:** Schematic of experiments. Human AC16 cells undergo 24 hours of thapsigargin or tunicamycin-induced ER stress, upon which cells are harvested. Total peptides and mPBA-enriched glycopeptides are analyzed using mass spectrometry. **B:** Canonical UPR markers are robustly induced with both thapsigargin and tunicamycin. Box: IQR; whiskers: 1.5x IQR. **: P < 0.01, ***: P < 0.005, ****: P < 0.001, two-tailed t-test (n=5). **C:** PCA plot of AC16 samples using the top 250 most variable glycopeptidoforms. **D:** Scatterplot of the fold-change correlation between protein fold change on the x-axis and the median glycopeptide-based fold change between thapsigargin and control (left) and glycopeptidoform on the (center). Bar plot depicting the maximum range of fold change values based of glycopeptidoforms compared to proteins (right). **E:** Volcano plot of showing differential abundance of glycopeptidoforms after thapsigargin treatment. x-axis: log2 fold-change; y-axis: –log 10 adjusted P values. **F:** As in E, but for AC16 cells after 24 hours of tunicamycin treatment. **G:** Mechanism of action of thapsigargin and tunicamycin.

Using an optimized condition (pH 10.5, 60% ACN, 0 mM KCl), we next evaluated the effect of loading various total masses of human HeLa total protein digest, ranging from 100 µg to 1 mg total peptide input. The average number of glycopeptide spectrum matches (gPSM) ranged from 7715 to 17233 across the input masses (**Figure 1H**). Because the gPSMs may map to indistinguishable species, we counted the number of glycosylation sites and distinguishable structural glycan combinations. We introduce the term glycopeptidoform here to refer to each unique instance of combined glycosylation site and glycan heterogeneity that is mappable by mass spectral evidence, in contradistinction to the term glycoform which may refer to the glycan portion micro-heterogeneity alone regardless of the site it is attached to, or the term glycopeptide, which may be used to refer to deglycosylated peptides regardless of the attached glycan in deglycoproteomics experiments (13). This terminology also provides a clear parallelism to the glycoproteoform concept, which refers to the total micro-heterogeneity and macro-hetergeneity across multiple glycosylation sites within a protein molecule. In total, we find that the number of unique glycopeptidoforms plateaus at 7916 at 500 µg input (**Figure 1H**). A considerable coverage remains detectable at 7715 gPSM and 3879 glycopeptidoform from as little as 100 µg total input peptides (**Figure 1H**), producing thousands of confident gPSM with manually verifiable spectral evidence, whereas the number of identifiable glycopeptidoforms did not further increase from 500 µg to 1 mg input, suggesting we are nearing saturation with regard to the sensitivity of our mass spectrometry system. We further measured the specificity of glycocapture by considering the proportion of gPSM vs. non-glycosylated PSM. At 500 µg input, specificity is at 77–94% suggesting most of the eluent peptides are glycosylated. Finally, duplicate glycocapture experiments from independent biological replicates show high correlations in glycopeptidoform intensity (R^2^: 0.667 to 0.835) across the input concentration range (**Supplemental Figure S1**). Taken together, these results show that mPBA allows highly sensitive enrichment and identification of intact glycopeptides (up to ∼8,000 glycopeptidoforms per injection) at low input (0.1 to 0.5 mg) without requiring further pre-fractionation.

### ER stress induces broad N-glycoproteome remodeling independent of protein expression

Next, to evaluate the effect on glycoproteostatic stressors on the human N-glycoproteome leading to ER stress, we examined ER stress models in the human AC16 cell line, chosen in part for its ease of expansion and its hybrid myocyte and fibroblast origin. AC16 cells abundantly express large muscle and extracellular matrix proteins and have been widely used to study ER stress response under thapsigargin and tunicamycin treatments (37, 38). The UPR pathway undergoes several distinct phases of responses following intrinsic timers, comprising an initial suppression of translation via the PERK-EIF2A axis within ∼1 hour, and transcriptional programs driven by ATF4, ATF6 and IRE1a/XBP1s after 2 to 8 hours (10, 39, 40). Notably, post-UPR proteomic changes are known to lag behind the acute-phase transcriptome and translatome remodeling, and may require up to 16–24 hours to catch up with transcriptional programs (40). We therefore opted to compare total peptides and intact glycopeptides after 24 hours of stressors to evaluate the proteome and glycoproteome response at new post-stress translational and post-translational set points (**Figure 2A**). Upon 24-hour treatment with 1 μM thapsigargin and 1 μg/mL tunicamycin, ER stress and UPR markers are robustly induced in AC16 cells, including the major ER Hsp70-family chaperone binding immunoglobulin protein/78 kDa glucose-regulated protein (BiP/Grp78, also known as HSPA5), ER chaperone and BiP nucleotide exchange factor Grp170 (also known as hypoxia up-regulated 1 HYOU1), ER protein folding factor protein disulfide isomerase 1 (P4HB/PDIA1), and ER Hsp90 chaperone 94 kDa glucose-regulated protein (Grp94/HSP90B1; also known as endoplasmin) (t-test P < 0.001) (**Figure 2B**). Proteome-wide quantification of over 6,000 proteins in each condition further confirms a robust proteome-wide response. Notably, despite their different ER stress induction mechanisms, thapsigargin and tunicamycin lead to highly correlated protein level log fold-changes (R^2^= 0.57; r=0.75) indicating overall similar total proteome profiles upon UPR activation (**Figure S1B**). This observation remains true on a pathway level, where both stressors induce UPR-related genes enriched in “PERK regulates gene expression”, “Translation”, “ATF4 activates genes in response to endoplasmic reticulum stress”, “IRE1alpha activates chaperones” and other Reactome terms (GSEA FDR-adjusted P ≤ 0.01) (**Supplemental Figure S2**). Therefore, the stressors robustly induce proteome signatures of ER stress and UPR, with clear concordance of fold-change in both specific UPR markers and across the proteome.

To quantify the parallel N-glycoproteomic changes, we next applied the mPBA workflow to determine the effect of these stressors on intact glycopeptides, analyzing a median of 2797, 3096, and 1524 glycopeptidoforms (total 4129 glycopeptidoforms) from 4675, 4956, 2347 gPSM across control, thapsigargin, and tunicamycin cells, respectively (after removing one outlier due to lack of signals, n=5 for control and tunicamycin; n=4 for thapsigargin) (**Supplemental Figure S3**). Notably, the top 250 most variable glycopeptidoforms recapitulate the sample group separation in the first two principal components, indicating glycosylation status alone is sufficient to distinguish the cellular stress and drug treatment status (**Figure 2C**). We first focused on the glycosylation ratios between thapsigargin and control cells. Importantly, only a modest correlation exists between protein fold-change and glycosylation fold-changes, measured as the average glycopeptidoform ratios at each glycosylation site, upon thapsigargin treatment (R^2^: 0.17). This correlation became even lower when we considered individual glycopeptidoform intensities were considered (R^2^: 0.13) (**Figure 2D**). This result indicates that glycoform provides orthogonal information to the cells than protein up-down regulations. In other words, the measured intact glycopeptide changes are not simply a reflection of protein differential abundance. This observation is also corroborated by considering the maximal range of absolute fold-change between glycopeptidoforms and proteins. Whereas most protein abundance changes are within a 10-fold absolute difference to control cells (i.e., 1000%; **Figure 2D** center); the magnitudes of glycopeptidoform changes are considerably more dynamic than protein abundance, with some glycopeptidoforms being over 50-fold (5000+%) more abundant under ER stress than in normal control cells (**Figure 2D**, right). This result illustrates that N-glycosylation is, at least in part, independently modulated from polypeptide chain synthesis under ER stress conditions. And moreover, the quantification of glycosylation status may provide a greater dynamic range of information to sensitively assess protein functional state than total protein pool size

We next considered the identities of individual glycopeptidoform changes and contrasted the thapsigargin and tunicamycin responses. Thapsigargin is a small molecule inhibitor of the sarco/endoplasmic reticulum calcium ATPase (SERCA), leading to the depletion of ER luminal calcium. The depletion of calcium disrupts protein folding and further inhibits calcium-sensitive ER glycoproteostatic machineries including UGGT1 and the lectin chaperones CNX and CRT. In contrast, tunicamycin, inhibits DP-GlcNAc:dolichol phosphate N-acetylglucosamine-1-phosphate transferase (DPAGT1). Inhibition of DPAGT1 prevents the synthesis of the N-glycans leading to a global arrest of N-glycosylation. Tunicamycin treatment led to substantial fewer unique glycopeptides and gPSM, consistent with its mechanism to globally abrogate the initial steps of protein N-glycosylation (**Supplemental Figure S3**). This is corroborated at an individual glycopeptidoform level, where we observe robust induction of multiple glycopeptidoforms upon thapsigargin including in Grp170/HYOU1, TSP1, and LYOX (**Figure 2E; Supplemental Data S1**). In contrast, far fewer instances of increased glycosylation is observed in tunicamycin (**Figure 2F**), consistent with the differential mechanisms of the two ER stress induction agents (**Figure 2G; Supplemental Data S1**). This result provides a striking contrast with the high concordance of the cellular response to the two stressors at the proteomic level (**Supplemental Figure S2**). Therefore, the observed glycosylation induction upon thapsigargin likely represents the outcomes of UPR-led production of new proteins and N-glycosylation, as the tunicamycin-led abrogation of total N-glycosylation renders cells unable to mount an overall glycome remodeling despite UPR activation and total proteomic response.

To further examine the how the observed glycosylation changes influence glycoproteins, we focused on select glycoproteins with multiple differentially regulated glycopeptidoforms. Grp170/HYOU1 is a key protein in remediating protein folding upon UPR, by acting as an UPR-induced chaperone as well as a nucleotide exchange factor to accelerate the BiP/Grp78 chaperone cycle. Grp170 is decorated by multiple N-glycosylation glycans at multiple sites. Inspection of individual protein changes shows that thapsigargin induces a number of specific glycopeptidoforms, including Grp170 HexNAc_2_Hex_9_ at N931 (logFC: 5.2; P.adj: 3.3e–4), N596 (logFC: 3.4; P.adj: 8.4e–3), and N515 (logFC: 5.3; P.adj: 4.5e–3) but to different extents, indicating significant macro-heterogeneity of differential glycosylation within the same protein under stress. Importantly, the magnitudes of some of these changes exceed that of the protein level in Grp170. This corroborates that in addition to the overall expression of Grp170 being modulated by the UPR via the IRE1/XBP1s and ATF6 arms (41), Grp170 also becomes relatively more glycosylated on a per-molecule basis. In contrast, we observe a decrease in glycosylation in FKB10 N352 HexNAc_2_Hex_6_ (logFC: –2.9; P.adj: 5.7e–4), RCN3 N140 HexNAc_2_Hex_6_ (logFC: –5.6; P.adj: 3.4e–4), and other glycopeptidoforms. Thus, ER stress alters the N-glycoproteome in a protein-, site-, and glycan-specific manner, reflecting regulations over protein abundance changes.

### Glycan classes differ broadly across proteins, sites, and localizations

Taking advantage of the glycan-peptide connectivity of the data, we assessed the effect of glycoproteostatic disruptions on secretory pathway compartments. Using UniProt annotations on protein localization (29) and cardiac cell surface marker data (42), we compared the N-glycosylation profiles of select ER/Golgi resident, lysosome, and cell surface proteins (**Figure 3A**). ER proteins including Grp94 and Grp170, RCN1, and CALU are modified primarily with high-mannose glycoforms with few sialylated and fucosylated forms detected, consistent with these proteins being primarily exposed to early glycan processing steps in the ER prior to hybrid/complex glycan processing (**Figure 3B**). In contrast, the lysosome membrane proteins LAMP1 and LAMP2 feature pervasive hybrid/complex glycoforms including fucosylated and sialylated mature glycans. The cell surface membrane proteins BSG and SLC3A2 feature a number of sialylated glycoforms. Extracellular matrix proteins such as COL5A1 and LAMB1 and LAMC1 feature relatively few sialylated glycoforms. Thus, the intact glycopeptide data readily recapitulate differences among classes of glycans based on the subcellular localization of the proteins.

**Figure 3:**
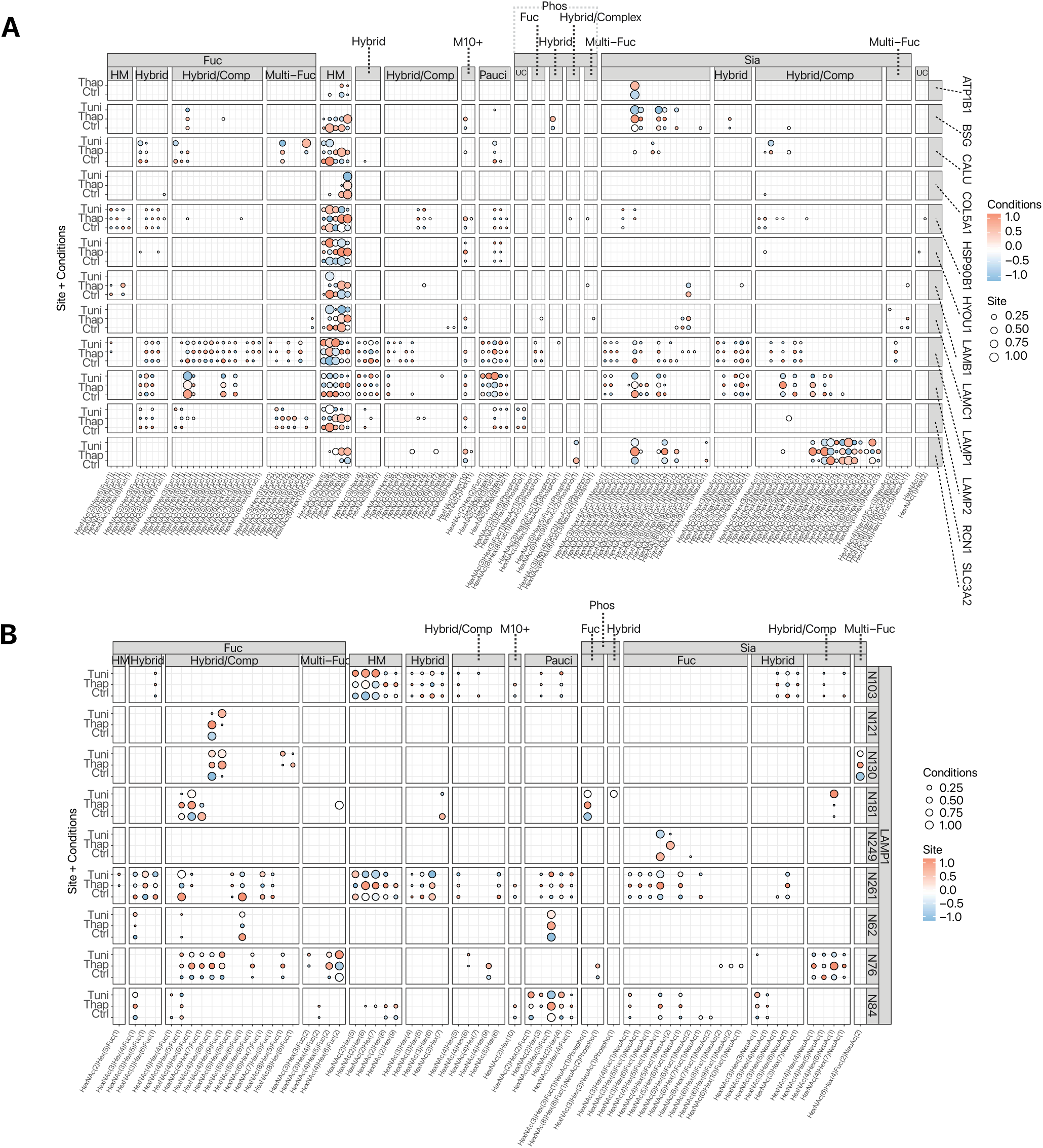
Glycoproteome remodeling broadly affects cell surface and intracellular proteins. **A:** Glycan changes in select ER/Golgi, lysosome, and cell surface proteins. Each data point represents one glycoform within each protein. The glycoforms (x-axis) are grouped by their glycan classes (top). Data point size: normalized relative abundance per protein per treatment condition. Color: Standardized relative abundance per glycoform across normal, thapsigargin, and tunicamycin treatment conditions. **B:** Glycopeptidoform changes in LAMP1. Each data point represents one glycopeptidoform (glycoform within a glycosite) within LAMP1. The glycoforms (x-axis) are grouped by their glycan classes (top). Each row facet is a glycosylation site within LAMP1. Data point size: normalized relative abundance per site per treatment condition. Color: Standardized relative abundance per LAMP1 glycopeptidoform across normal, thapsigargin, and tunicamycin treatment conditions.

Focusing on LAMP1, the data reveals a complex N-glycosylation pattern with 301 total glycopeptidoforms across 9 glycosylation sites. This coverage is comparable to that of a prior work that performed targeted profiling on immunoprecipitated LAMP1 (43), highlighting the sensitivity of the mPBA approach to provide comprehensive glycan information across proteins. The data portrays an interaction between glycosylation sites and glycoforms in LAMP1 (**Figure 3C**). For instance, the N103 site features largely high-mannose glycans and very few hybrid/complex glycoforms, whereas the N261 site contains a large variety of both high-mannose and hybrid/complex glycoforms, the N76 site features mostly fucosylated or sialylated hybrid/complex glycoforms, and the N84 site features largely paucimannose forms. Interestingly, thapsigargin leads to an increased abundance of high-mannose form in the N261 site but not the N103 site. Thus, the mPBA data reveals that within one protein, different N-glycosylation sites not only feature different glycan classes but these classes can alter in response to stress in different manners.

### Stress induced N-glycosylation changes are consistent with specific protein maturation and trafficking defects

To gain deeper insights into the individual regulation of glycosylation events under stress, we examined two proteins with substantial micro-heterogeneity and macro-heterogeneity in glycosylation profiles, focusing on Grp94 and laminin subunit gamma-1 (LAMC1). Grp94 (HSP90B1; endoplasmin) is a stress-inducible ER-resident chaperone of the Hsp90 family, with a specific client list that includes multiple cell surface localized and secreted proteins. There is contrasting information on the glycosylation of Grp94 in the literature. Under baseline conditions, Grp94 is thought to be predominantly monoglycosylated at one site (N217) whereas “hyper-glycosylation” of Grp94, at the five facultative sites N62, N107, N445, N481, and N502, is assumed to occur only during stress or cancer related Grp94 overexpression (44). In the experiment, we identified glycopeptidoforms at all 6 known Grp94 glycosylation sites (N62, N107, N217, N445, N481, and N502). The N217 site, corresponding to the constitutive mono-glycosylated form of Grp94, is associated with the greatest number of attached glycoforms (**Figure 4A**). On the other hand, N107, N445, N481, and N502 are modestly glycosylated, consistent with these facultative sites being minimally modified in the baseline. Surprisingly, we find a considerable degree of N62 glycosylation even under baseline conditions, suggesting some hyperglycosylated Grp94 exists in unstressed AC16 cells.

**Figure 4:**
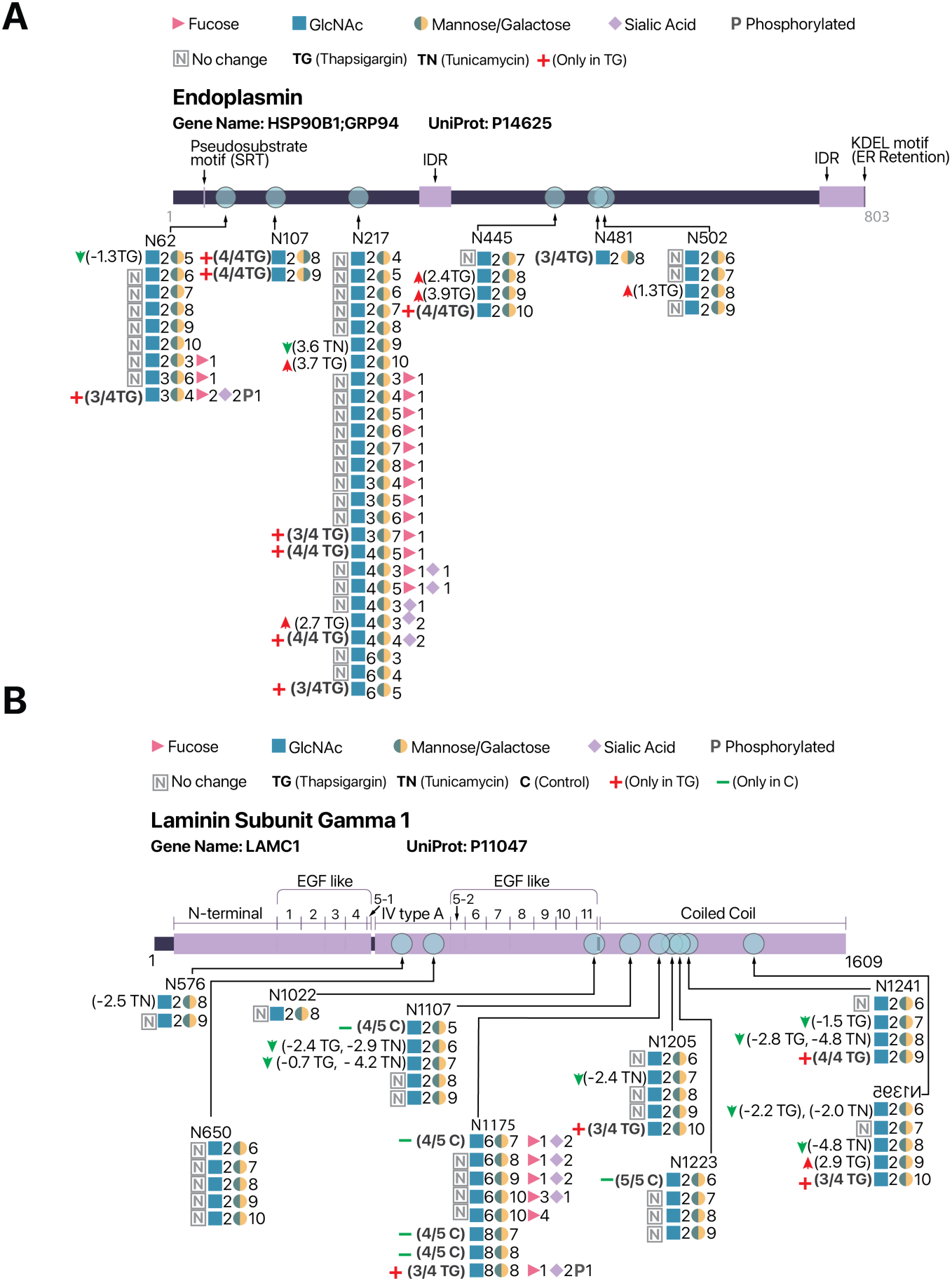
ER stress invokes protein-, site-, and glycan- specific N-glycosylation remodeling. **A:** Map of quantified glycopeptidoforms along the Grp94 (HSP90B1) protein sequence, showing the heterogeneity of glycosylation both across N sequons and across glycans within each site. (…) **B:** As in A, but for laminin subunit gamma 1 (LAMC1)

The glycosylation of N62 within the N terminal unstructured region is previously shown to be necessary for Grp94 cell surface translocation (8), consistent with the existence of a non-ER pool in unstressed cells. The possibility of a cell surface pool is further supported by the micro-heterogeneity of attached glycan structures, where we identified over 20 fucosylated, sialylated, or complex glycoforms. This extensive number of glycoforms that are modified by Golgi glycosyltransferase and glycosidases strongly indicates that although possessing the ER retention KDEL sequence and canonically annotated as being mostly an ER resident protein, endogenous Grp94 undergoes prolonged Golgi residency or exists with substantial non-ER pools even under baseline conditions.

Upon ER stress induction, a subset of glycopeptidoforms show varying degrees of differential abundance, including the gain of 7 new glycoforms across the known glycosylation sites, parallel to the increased abundance of 4 glycoforms and decreased abundance of 1 glycoform in stressed cells, highlighting a complex remodeling of glycosylation status. At the constitutive N217 site, we find an increase in glycosylation of HexNAc_2_Hex_10_, which is consistent with CNX/CRT binding and/or reglucosylation (GlcNAc_2_Man_9_Glu_1_) by UGGT1, indicating the induction of Grp94 is accompanied by prolonged folding and refolding cycles in the ER. The increase in glycoform abundance at the N445 and N502 site is likewise consistent with hyperglycosylation of Grp94 under increased expression level. At the N62 site, we find that thapsigargin led to the gain of a HexNAc_3_Hex_4_Fuc_2_Sia_2_P_1_ form that corresponds to the mannose-6-phosphate signal for lysosome targeting. At the same time, there is a gain of multiple new glycoforms including complex, fucosylated, and sialylated glycoforms, consistent with increased Golgi passage and a possible increase in the relative size of the cell surface localized epichaperone pool. Thus, the glycopeptidoforms of Grp94 portrait a complex picture of its regulation by glycosylation, with evidence of increased aberrant glycosylation of a subpopulation of unstructured Grp94 protein (N445 glycosylation).

We next examined laminin subunit gamma-1 (LAMC1) as a second exemple protein with highly heterogeneous glycosylation sites and glycoforms (**Figure 4B**). Laminins are a major component of the basement membrane and are composed of heterotrimers of alpha, beta, and gamma subunits. Cardiac cells are thought to express mostly laminin-221, which are heterotrimers of the alpha-2, beta-2, and gamma-1 isoforms. The data reveals a wide diversity of 39 glycopeptidoforms LAMC1 in AC16 cells, distributed broadly across 9 known N-glycosylation sites (N576, N650, N1022, N1107, N1175, N1205, N1223, N1241, N1395) as annotated on UniProt. Notably, ER stress induced glycosylation changes in LAMC1 are preferentially found among N-sequons located towards the C-terminus coiled coil domain (N1107, N1175, N1205, N1223, N1241, N1395) (**Figure 4B**). We observe a loss of complex glycans at the N1175 site in the coiled coil domain coupled to a reduction of Man6/Man7/Man8 forms (B1241 and N1395), pointing to decreased mannose trimming. These results point to a possible model of a delay in mannose trimming and anterograde transport defects during protein maturation, where the early steps of monomer folding may proceed normally but aberrant glycosylation occurs after the initial rapid phase of globular domain folding. Taken together, the analysis of glycopeptidoform quantification reveals both microheterogeneity and macro-heterogeneity of Grp94 and LAMC1 in a manner that maps to distinct processing steps and functional pools during the maturation process.

### Mannose trimming is a common defect across endomembrane proteins under ER stress

Following the results Grp94 and laminin subunit gamma-1, we asked whether global site-specific glycosylation changes in ER stress would also reflect general protein maturation dysfunctions. To discern the effect of glycosylation changes along the endomembrane pathway, we queried the glycan classes across identified glycopeptidoforms, by summarizing both the linked glycan types and the protein identities of the linked glycosylation sites. Strikingly, the summarized data reveal a strong accumulation of glycopeptides with high-mannose glycans upon ER stress, coupled to a decline of glycopeptides with fewer mannose subunits (**Figure 5A**). Specifically, grouping of MSFragger-Glyco glycan assignments across intact glycopeptides shows a statistically significant increase in HexNAc_2_Hex_10_ as well as HexNAc_2_Hex_9_ intensities in thapsigargin. The former glycan corresponds to the GlcNAc_2_Man_9_Glc_1_ form in the ER, which occupies an early critical step in protein maturation. As mentioned, newly glycosylated proteins are deglucosylated by α-glucosidases I and II to create monoglycosylated GlcNAc_2_Man_9_Glc_1_ and where the protein becomes competent for binding the major ER lectin chaperons calnexin and calreticulin for folding. If successfully folded, the GlcNAc_2_Man_9_Glc_1_ modification is further trimmed to GlcNAc_2_Man_9_, corresponding to the latter HexNAc_2_Hex_9_, that is competent for anterograde transport, whereas unfolded proteins may be reglucosylated by UGGT1 for additional opportunities to fold. As the high-mannose glycans occur early in the protein glycosylation and maturation pathways, the accumulation of HexNAc_2_Hex_10_ and HexNAc_2_Hex_9_ indicates prolonged UGGT check point cycle and reduced folding capacity. At the same time, there is a significant decrease in the overall intensity HexNAc_2_Hex_6,_ which reflects a glycoform that has undergone further trimming of mannose units, either in the ER through EDEM1/3 for unfolded proteins, or in the Golgi by alpha mannosidases MAN1A1, MAN1A2. The observed depletion of trimmed glycans in turn indicates that under thapsigargin, there is a reduced ER-to-Golgi transport of properly exported proteins, and/or a deficiency of EDEM capacity prior to ERAD.

**Figure 5:**
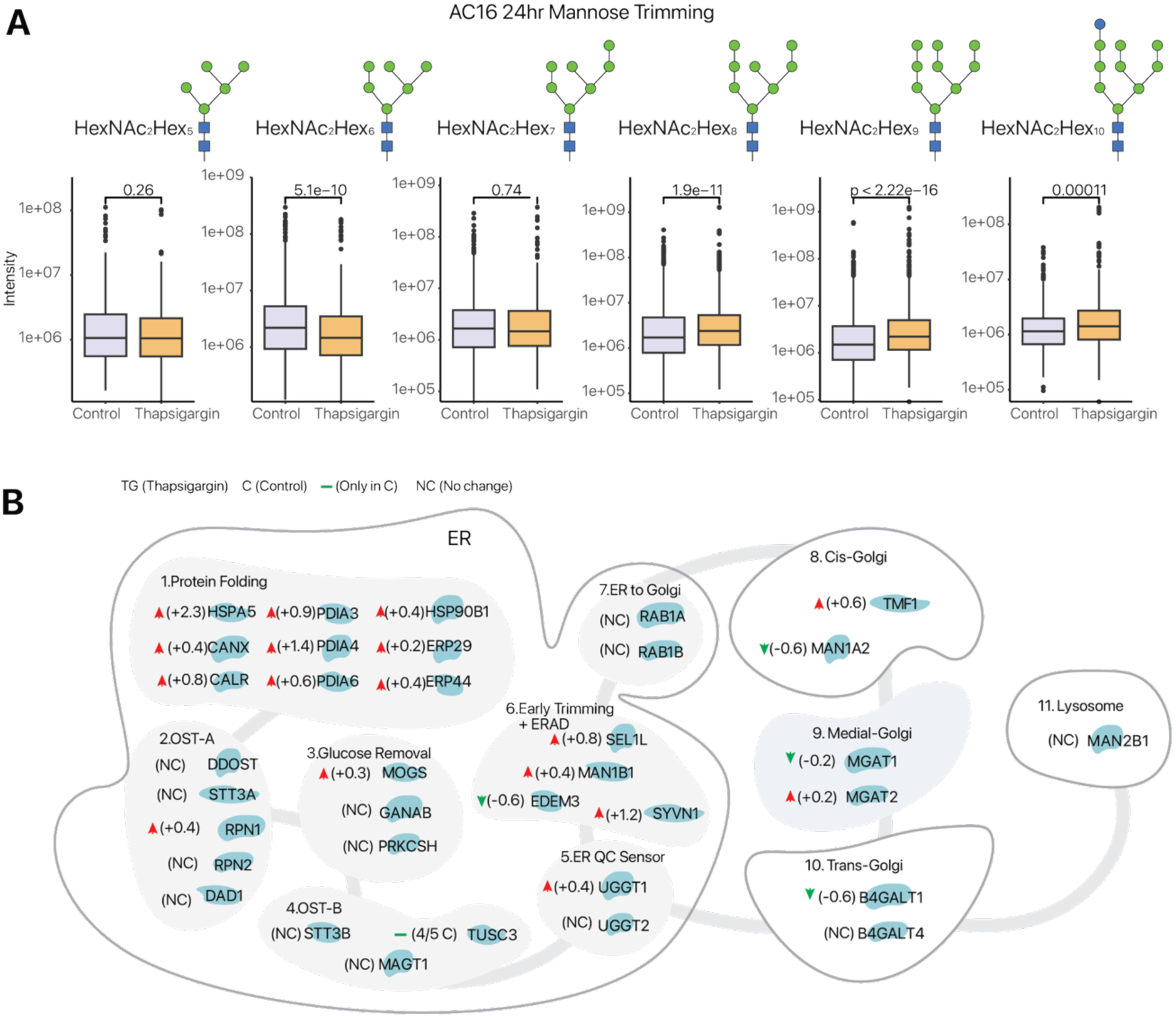
Mannose trimming defects are a conserved lesion across secretory pathway proteins under ER stress. **A:** Boxplots detailing alterations in glycoforms in the mannose trimming pathway with thapsigargin treatment. Each panel represents a specific N-glycan during the trimming process using common glycopeptides with at least 2 non-zero values. Box: IQR; whiskers: 1.5x IQR. P values: two-tailed t-test. **B:** Total protein abundance changes of key ER chaperones and glycogenes along the secretory pathways. Numbers denote log2 foldchange in protein level in thapsigargin over normal cells. NC: no significant change.

To examine the potential causes of high-mannose glycan accumulation, we measured the effect of thapsigargin on total protein abundance of critical chaperones, glycosyltransferase, and other resident ER proteins involved in protein folding, transport, glycosylation, and glycan processing (**Figure 5B**). As expected, we find a uniform increase in major UPR-induced ER chaperones, which provides validation to the proteomics data. Notably, we find UGGT1 protein level to be significantly elevated, consistent with up-regulation of UGGT checkpoint refolding/reglucosylation cycles. In parallel, we find a complete loss of detectable TUSC3, a tumor suppressor protein with still poorly defined role, but which recently has been shown to be part of the OST complex that functions to alter early glycan attachment to specific proteins (45). At the same time, there is a significant decrease in the ER ERAD mannosidase EDEM3, responsible for the trimming of GlcNAc_2_Man_8_ on unfolded proteins to GlcNAc_2_Man_7_, the latter of which exposes the ɑ1,6 mannose linkage that is recognized by ERAD lectins to commit the proteins for degradation. This is further accompanied by a significant decrease of MAN1A2, a Golgi-resident ɑ-mannosidase that functions in glycan maturation prior to the formation of complex glycans, pointing to a mechanism for the global accumulation of high-mannose and depletion of trimmed-mannose glycopeptidoforms.

Finally, to confirm whether the observed glycoproteome remodeling and mannose trimming defects are conserved across cell types, we applied the mPBA workflow toward analyzing human induced pluripotent stem cell-derived cardiomyocyte (hiPSC-CM) (**Figure 6A**). Intact glycopeptide analysis following mPBA identified a total of 4777 glycopeptidoforms across all normal and stress conditions from 0.2 mg input per sample, indicating mPBA also allows sensitive glycoproteome analysis in differentiated cell types. As hiPSC-CM are post-mitotic cells, they have a different proteostatic requirements as reflected by a reduced protein synthesis load under longer median protein half-life than AC16 cells (37). Consistently, the glycoproteome and proteome of hiPSC-CM continued to evolve under ER stress past the first 24 hours of stress induction, with ER stress response proteins BiP/Grp78, Grp170, and Grp94 only reaching a similar level of induction as AC16 cells after 72 hours (**Figure 6B; Supplemental Figure S4**). We therefore first examined the glycopeptidoform changes of hiPSC-CM at 72 hours of treatment. Strikingly, mannose trimming showed a similar pattern as AC16, suggesting global mannose trimming defect is a conserved feature of thapsigargin-induced ER stress between two cell types (**Figure 6C**). To further investigate the temporal dynamics, we considered also the glycoproteomics data acquired at an earlier time point (24 hours) post-thapsigargin treatment. Despite relatively modest induction of UPR-regulated proteins at this point, the accumulation of HexNAc_2_Hex_8–10_ and the depletion of HexNAc_2_Hex_5–6_ is already apparent, suggesting mannose trimming defects may precede further ER stress and UPR molecular sequelae (**Figure 6D**). Taken together, the glycoenzyme proteomics data in AC16 corroborate the intact glycopeptides data in AC16 and hiPSC-CM, pointing to mannose trimming as a critical lesion in the N-glycosylation remodeling of stressed cells.

**Figure 6:**
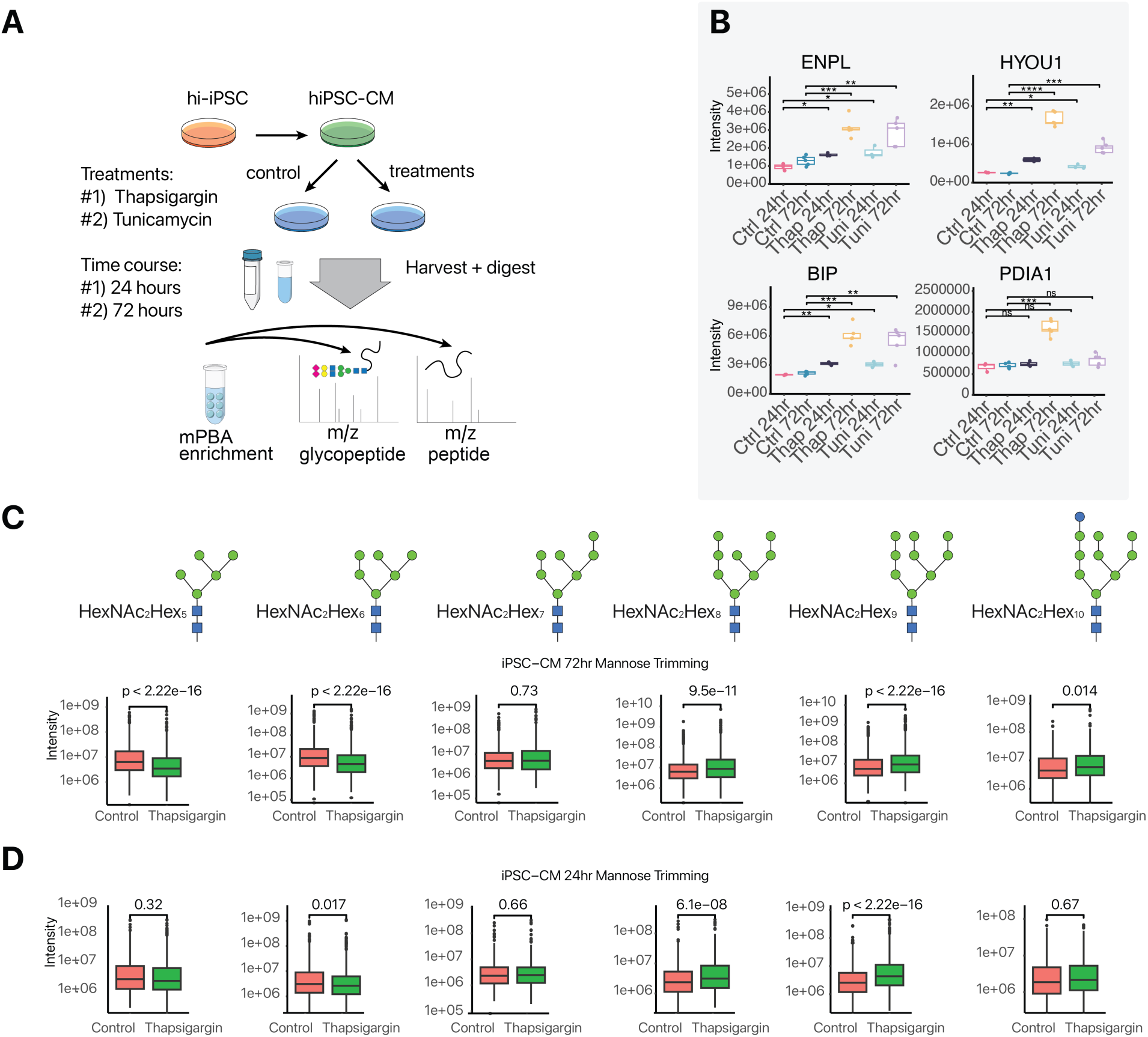
Mannose trimming defects are conserved in hiPSC-CM under ER stress. **A:** Experimental schematic of hiPSC-CM thapsigargin/tunicamycin treatment, and mPBA enrichment. **B:** Markers of ER stress of hiPSC-CM 24 and 72 hours. **C:** Evidence for mannose trimming defect in hiPSC-CM after 72 hours of thapsigargin treatment. Box: IQR; whiskers: 1.5x IQR. P values: two-tailed t-test. D. As in C, but for hiPSC-CM after 24 hours of thapsigargin treatment.

## Discussion

We describe mPBA as a method to capture and enrich multiple glycan classes for intact glycopeptide mass spectrometry analysis. Protein N-glycosylation is one of the most common and complex protein modifications (46–49) and is especially enriched among cell surface proteins that present common drug targets, but analysis of intact glycopeptide that preserves the connectivity between the protein (peptide) – glycan bond remains challenging due to their low abundance and inefficient ionization. Effective enrichment strategies of intact glycopeptides from complex mixtures therefore remain needed that can enable routine quantitative analysis between experimental conditions. To date, dozens of enrichment strategies have been employed using a wide variety of different chemistries bind glycopeptides (22, 50). Recent work using HILIC, ERLIC, and boronic acid enrichment for intact glycopeptide quantification by HCD have demonstrate the feasibility of scalable intact glycopeptide analysis from 2–4 mg of input total proteins/peptides (16, 17). Glycocapture by boronic acid chemistry occurs via a change in hybridization in the boron atom from sp^2^ to sp^3^ that results in the trigonal planer bonding arrangement to change to tetrahedral under basic conditions. In acidic conditions, hybridization is reversed resulting in the release of bound glycans. The formation of pH-reversible covalent bonds to cis-diol groups within a glycan should allow for a simple bind-and-release model for enrichment, but boronic acid-based enrichment has been relatively under-realized, in part due to the low binding strength. Recent works from multiple groups have shown that this can be remedied by boronic acid networks based on dendrimers or polymer meshes. Building on these advances, our implementation and optimization results show that glycan enrichment from 0.1–0.5 mg total peptide input is sufficient to support the confident identification and quantification of thousands of glycopeptidoforms on a common and accessible mass spectrometry platform. We foresee the optimization demonstrated here should help enable sensitive and quantitative comparisons of glycoprotein micro- and macro-heterogeneity cross cell states and experimental conditions.

Glycan class bias presents another important consideration for enrichment strategies. An effective glycopeptide isolation technique should be amenable to common human glycosylation structures and survey glycoproteins in both the endomembrane pathways as well as cell surface and/or extracellular space. Extraction methods using lectins is subject to the substrate recognition selectivity of the specific lectin being used. For instance, lectin phase extraction commonly employ concanavalin A, which preferentially captures high-mannose glycans that are more prominently found in proteins during early processing steps but are depleted in mature glycoproteins (15, 51). As boronic acid chemistry captures glycan containing 1,2- and 1,3-cis-diols that are found in virtually all mammalian glycans, it is potentially suitable for the enrichment of multiple glycan classes (17, 22, 36, 52). Despite boronic acid chemistry having been referred to as capturing glycoproteins universally in the literature (35, 36) however, its relative efficiency of capturing different glycan classes has not been systematically investigated. The optimization results here provide a framework to compare glycocapture efficiencies among glycan classes that may be differentially important for probing the status of mature and cell surface proteins. Our results show that different organic strengths may be suitable for biasing the capture of glycan classes based on experimental needs. By tuning solvent organic strengths, individually refined protocols may be possible. For instance, 50–60% ACN may be used to maximize total identification, whereas higher organic strengths may be employed for cell surface sialylated N-glycoproteome analysis, or low ACN% conditions when intracellular trafficking pathways are the focus.

Mutations in glycosylation enzymes underlie many congenital diseases of glycosylation, whereas aberrant N-glycosylation patterns often accompany in common age-associated diseases including cancer and neurodegenerations. Importantly, because the final glycan is not determined directly by a genome template but arise from the summed output of multiple orchestrated enzymatic sequences, a disruption in the capacity, specificity, and kinetics of processing steps could lead to a system failure in glycan processing, preventing the generation of the matured glycoforms (i.e., complex, fucosylated, and sialylated glycans) critical for the trafficking and function of endomembrane or cell surface proteins. Applying mPBA to interrogate the ER stress-induced remodeling of the N-glycoproteome, we observed a highly dynamic and heterogeneous N-glycoproteome, where ER stressors lead to highly variable protein-, site-, glycan-, and stressor-specific changes in numerous proteins. Strikingly, although thapsigargin and tunicamycin have similar effect on the UPR marker and proteome level, they show highly discordant glycopeptidome changes, indicating these commonly used ER stressors do not activate a uniform UPR program but may rewire the glycosylation code used in protein folding and trafficking. Under thapsigargin treatment, N-glycosylation remodeling at the glycopeptidoform level shows distinctive patterns with respective to known protein domains or folding steps. The data points to specific processing and maturation defects of endogenous proteins under ER stress, as illustrated in Grp94 and laminin subunit gamma-1.

Grp94 is a major chaperone of the Hsp90 family that can function alone in protein folding or in tandem with Hsp70. Grp94 is conventionally taken to be primarily mono-glycosylated only at the N217 site in non-stressed tissues, whereas glycosylation of other sites is thought to occur only during elevated Grp94 expression. Because some of these facultative sites are buried in structured regions, the hyper-glycosylation of Grp94 involving these sites was suggested to reflect unfolded Grp94 marked for ERAD (44). Other studies however have shown that Grp94 N62 glycosylation marks it for translocation to the cell surface, where it can act as a scaffolding epichaperone to alter protein-protein interactions (8). As prior works largely relied on mutagenesis and electrophoretic mobility shifts to assess glycosylation status, the site-specific glycosylation profile of Grp94 remained to be elucidated. Our data reveals that a number of glycoforms can already be confidently identified at the facultative sites in baseline AC16 cells, including at N62. Moreover, the glycosylation sites are associated with multiple distinct glycoforms, including fucosylated and sialylated forms typically associated with Golgi and cell surface proteins at N217. Upon stress, the number of glycoforms further increases. This pattern suggests the existence of multiple Grp94 subcellular pools, including a potential cell surface localized Grp94 pool at baseline conditions. Moreover, the diversity of glycosylation at facultative sites is consistent with a high multiplicity of structural conformations of Grp94 required by its stress response roles as both a protein foldase and a holdase (53). Protein modifications in the Hsp90 family proteins fine-tune their ATPase activity, stability, and co-chaperone and client binding in a manner has been referred to as a “chaperone code”. The data here reveal that the molecular multiplicity of chaperone codes likely involves a previously under-appreciated level of N-glycosylation micro- and macro- heterogeneity.

Laminins are large (∼400–900 kDa) heterotrimeric glycoproteins that require proper folding and assembly during its secretion to the extracellular space, after which individual trimers further polymerize into a basement membrane lattice. Within each of the alpha, beta, and gamma subunits, there is a variable short-arm comprised on globular and EGF domains (∼residues 46 – 1030 in LAMC1), followed by an α-helical coiled coil in the long arm (residues 1038 – 1609 in LAMC1) that functions in the trimerization by intertwining the respective coiled coil of the alpha, beta, and gamma subunits together (54). Surprisingly given its importance in muscular dystrophy, cell adhesion, and pluripotency maintenance; relatively little is known about the assembly sequence of laminin trimer. The results here reveal a bimodal behavior between the short-arm localized N-glycosylation sites and those at the coiled coil, and that the stress-induced changes disrupt the maturation processes associated with trimerization domains. This suggests a model where upon LAMC1 translation, the N-terminal globular domains fold first in an initial phase, followed by a second phase of dimerization, trimer formation, and disulfide bridges at the coiled coil domain while the final trimer is assembled. This model is supported by early biochemistry work that shows the β and γ subunits can independently form a dimer prior to α subunit incorporation and secretion (55–57). Although further work is needed to confirm the relationship between protein site- and domain-specific profiles and their trafficking status, the data here strongly hint at the utility of intact glycopeptide analysis for probing the disrupted biogenesis and maturation of individual proteins during stress and disease conditions.

Consistent with this notion, our glycoproteome-wide analysis shows that ER stress is accompanied by an accumulation of GlcNAc_2_Man_8–10_ forms and the depletion of accumulation of GlcNAc_2_Man_5–6_ forms. The sequence of N-glycosylation is essential for protein folding, quality control, maturation, and trafficking through the endomembrane pathway (4, 58). Mannose trimming is essential for both the processing of successfully folded proteins and the degradation of misfolded proteins in the ER. In the former, GlcNAc_2_Man_9_ (or GlcNAc_2_Man_8_ depending on the localization of MAN1B1) is competent for anterograde Golgi transport, whereas in the cis-Golgi, the glycans require trimming to GlcNAc_2_Man_5_ to be used by MGAT1 for further construction of hybrid and complex type glycans.

Whereas for misfolded proteins, trimming by EDEM1/3 in the ER to GlcNAc_2_Man_7_ is required for the misfolded proteins to be recognized by ERAD lectins for degradation (59). One question is whether ER or Golgi mannosidase catalyzed processes, or both, is disrupted under stress. Because GlcNAc_2_Man_7_ is already competent for ERAD, we reason that the accumulation of GlcNAc_2_Man_5–6_ indicates Golgi mannose trimming is at least partially responsible. This is corroborated by the total protein abundance measurements showing both EDEM3 and MAN1A2 to be significantly down-regulated by thapsigargin. As the Golgi is critical for the continued maturation of complex N-glycosylation structures, Golgi mannosidases in particular may present targets for remediating cell surface remodeling accompanying ER stress. Moreover, we note that the data on intact glycopeptides in turn provides strong evidence that ER stress changes the activity of these enzymes, and not only expression level. We speculate that these coordinated glycoenzyme changes including EDEM3/MAN1A2/TUSC3 may represent a stress-specific glycan editing switch, which would play important parts in triaging the fate of glycoprotein lifetime and trafficking in stress and disease situations.

### Limitations of the Study

This study demonstrates a comparison of intact glycopeptide changes under stress conditions. Current intact glycopeptide analysis by stepped-HCD fragmentation mass spectrometry has several inherent limitations: the compositions of glycans are inferred from the masses of monosaccharide classes (e.g., hexose vs. glucose) based on prior knowledge of mammalian glycome; some glycan isomers have the same mass and cannot be resolved; linkage information between glycans is not measured; and assignment may be complicated by isobaric glycopeptides. In addition to these common limitations, the subcellular localizations of proteins in our analysis are based on prior profiling experiments or protein annotations (e.g., ER resident vs. cell surface proteins); distinguishing the intracellular pool of a surface targeted protein will require cell surface labeling techniques (13, 42). To delineate glycoproteoforms (i.e., combination of glycoform configurations across multiple sites or polypeptide chains within the same molecule or complex), a combination of glycopeptide analysis and native/top-down mass spectrometry is required. Despite these limitations, we acquired proteome-wide insights into the role of site-specific glycoforms in stress response by providing a deep resource of the identity and quantity of thousands of glycopeptidoforms in normal and stressed human cells for the first time, and expanding the scope of the known mammalian ER stress regulome. The findings suggest that intact glycopeptide analysis may help identify lesions of protein maturation and trafficking disruptions along secretory pathways. More generally, this resuls showcase the utility of the mPBA enrichment as a sensitive method to enable N-glycoproteomics investigations. The use of magnetic bead conjugation renders the mPBA workflow compatible with robotic platform automation in large-scale studies. We foresee this workflow will be useful for understanding glycosylation changes in individual glycoform granularity and aid in studies to connect glycoforms to function.

## Supplemental Figures

**Supplemental Figure S1.**
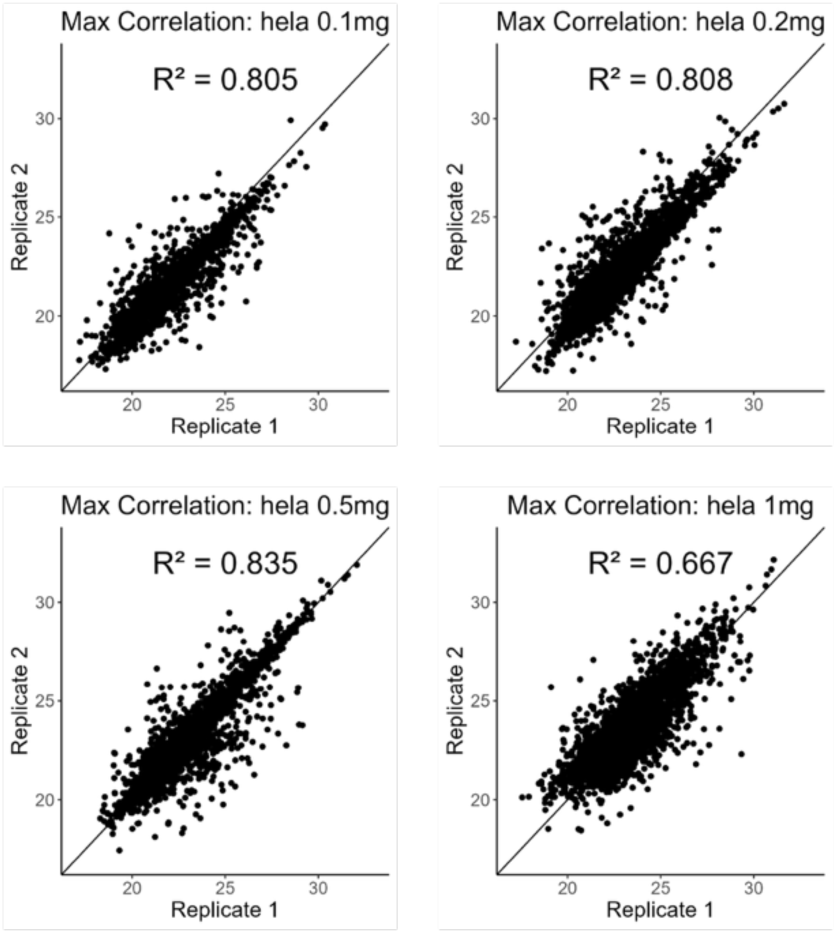
Correlation of mPBA enriched glycopeptide intensities across replicates and input amounts. Scatterplot represents MS1 intensities of glycopeptides across two biological replicates in each tested condition

**Supplemental Figure S2.**
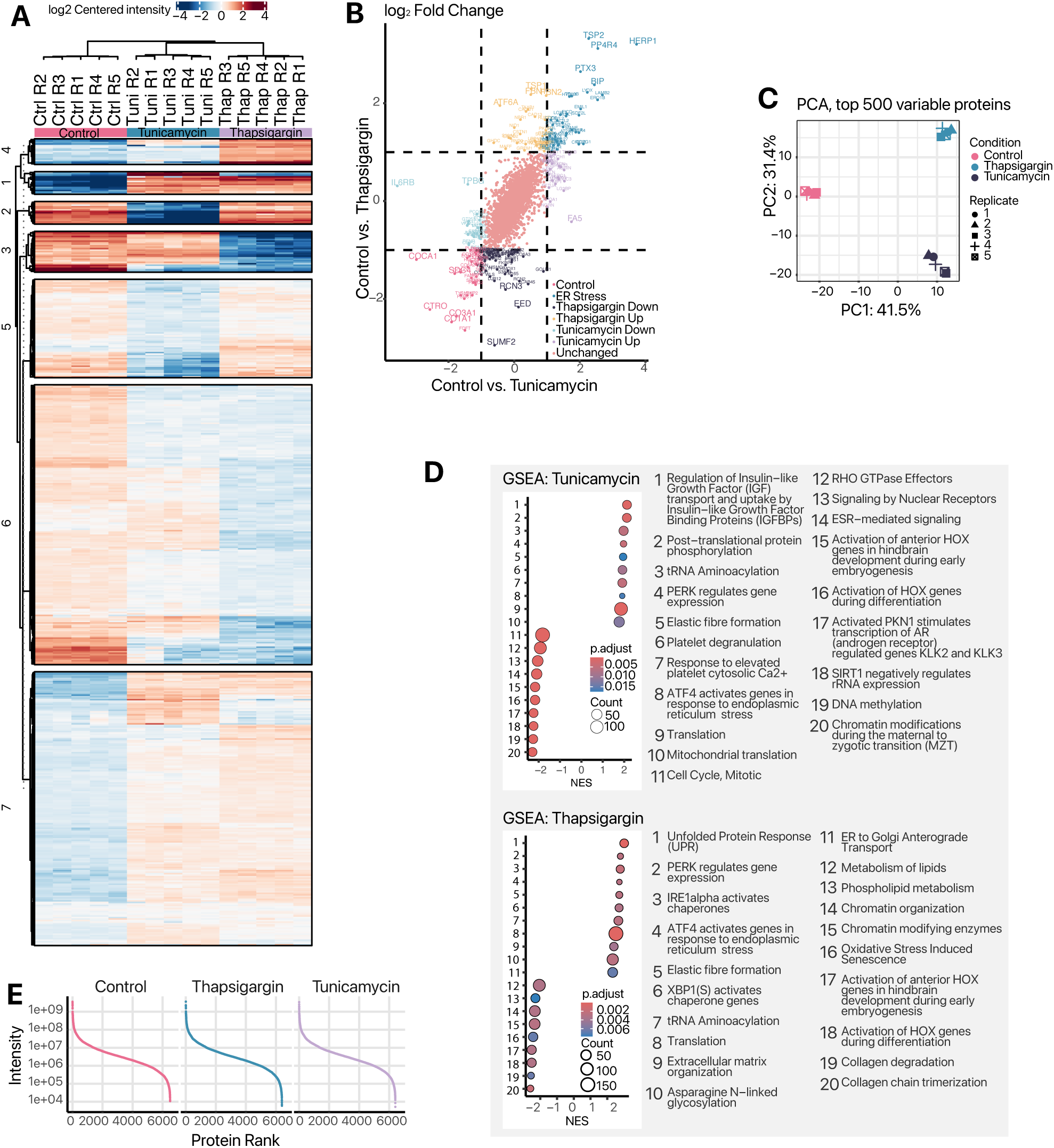
Thapsigargin and tunicamycin induce robust unfolded protein response in human AC16 cardiac cell. **A:** Heatmap of AC16 protein expression after exposure to 1 µM thapsigargin, 1 µg/mL of tunicamycin or DMSO vehicle for 24 hours. Color: log2 centered intensity; Numbers: clusters. **B:** Comparison of the log2-fold change between conditions. The fold change with thapsigargin is displayed on the y-axis and fold change with tunicamycin is on the x-axis. Proteins are colored based on their position in the matrix. Only proteins that have a p-value less than 1 are shown outside the “unchanged” box. **C:** Principal component analysis of protein abundance in normal, thapsigargin, and tunicamycin cells. **D:** Gene set enrichment analysis (GSEA) results using the Reactome pathway database to evaluate the top 10 most upregulated and top 10 most downregulated pathways upon exposure to thapsigargin (left) and tunicamycin (right). **E:** Dynamic range of protein LFQ intensity over protein rank in each experiment.

**Supplemental Figure S3.**
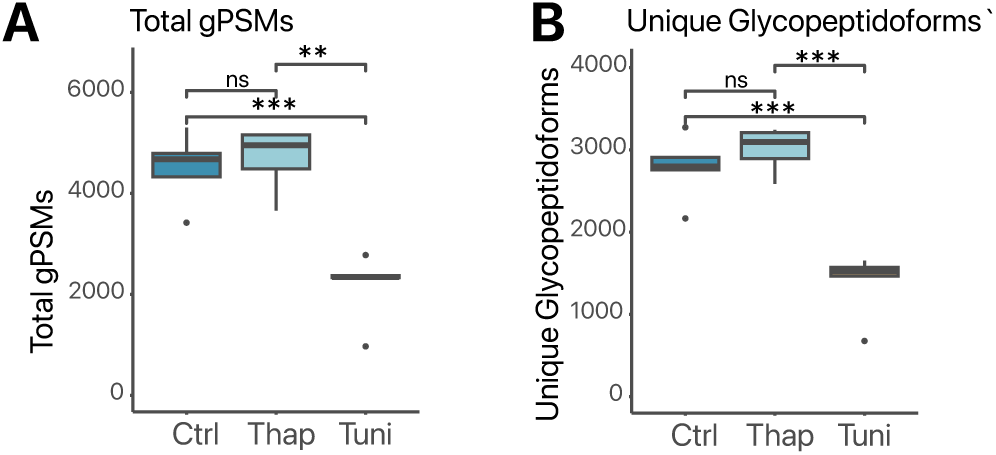
The effect of ER stress on the AC16 intact glycopeptide identification. **A:** Boxplots showcasing the total number total glycopeptide-spectrum matches (gPSMs) from AC16 cells after enrichment with the mPBA method. Box: IQR; whiskers: 1.5x IQR. *: P < 0.05; **: P < 0.01; ***: P < 0.005; ****: P < 0.001; ns: not significant; two-tailed t-test **B:** As in A, but for the number of unique glycopeptidoforms.

**Supplemental Figure S4.**
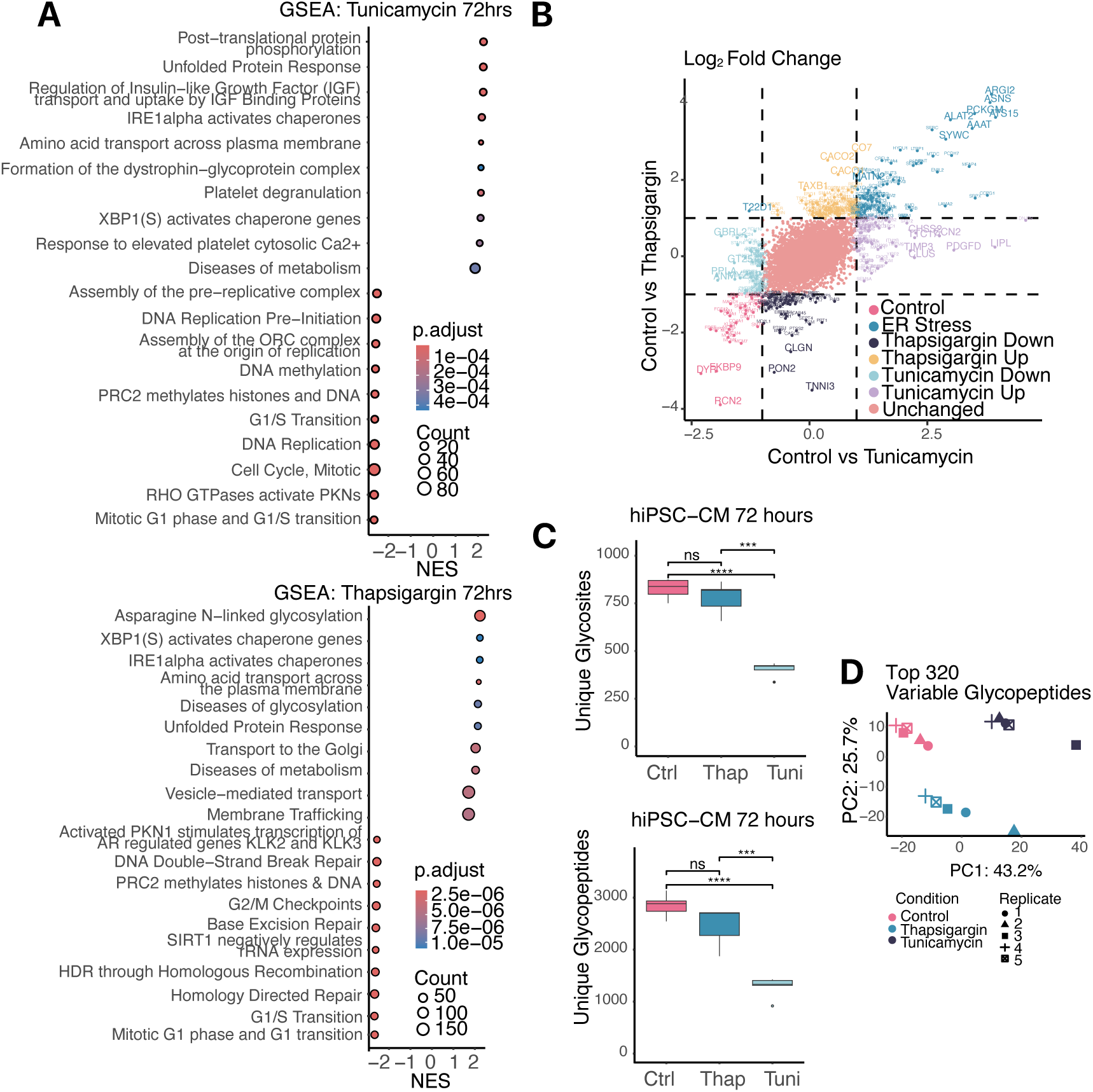
The effect of ER stress on the hiPSC-CM proteome and glycoproteome. **A:** GSEA dotplot results using the Reactome pathway database to evaluate the top 10 most upregulated and downregulated pathways upon exposure to 1 µM thapsigargin, 1 µg/mL of tunicamycin or DMSO vehicle for 72 hours. Size: gene count; Color: GSEA FDR-adjusted P value. **B:** Comparison of the log2-fold change between conditions at 72 hours. The fold change with thapsigargin is displayed on the y-axis and fold change with tunicamycin is on the x-axis. Proteins are colored based on their position in the matrix. **C:** Boxplots detailing the number of glycosites and unique glycopeptides identified in hiPSC-CM after 72 hours of treatment. Box: IQR; whiskers: 1.5x IQR; ***: P < 0.005; ****: P < 0.001; ns: not significant; two-tailed t-test. **D:** PCA plots of each of the glycoproteomic hiPSC-CM samples using the top 320 (72 hours) most variable glycopeptidoforms.

